# Alpha oscillations are involved in localizing touch on hand-held tools

**DOI:** 10.1101/2021.10.28.466268

**Authors:** Cécile Fabio, Romeo Salemme, Eric Koun, Alessandro Farnè, Luke E. Miller

**Affiliations:** Integrative Multisensory Perception Action & Cognition Team - ImpAct, Lyon Neuroscience Research Center, INSERM U1028, CNRS U5292, Lyon, France; University of Lyon 1, Lyon, France; Hospices Civils de Lyon, Neuro-immersion, Lyon, France; Center for Mind/Brain Sciences, University of Trento, Rovereto, Italy; Donders Institute for Brain, Cognition and Behaviour, Nijmegen, the Netherlands

**Keywords:** Tactile localization, tool use, extended sensing, alpha oscillations

## Abstract

The sense of touch is not restricted to the body but can also extend to external objects. When we use a hand-held tool to contact an object, we feel the touch on the tool and not in the hand holding the tool. The ability to perceive touch on a tool actually extends along its entire surface, allowing the user to accurately localize where it is touched similarly as they would on their body. While the neural mechanisms underlying the ability to localize touch on the body have been largely investigated, those allowing to localize touch on a tool are still unknown. We aimed to fill this gap by recording the EEG signal of participants while they localized tactile stimuli on a hand-held rod. We focused on oscillatory activity in the alpha (7-14 Hz) and beta (15-30 Hz) range, as they have been previously linked to distinct spatial codes used to localize touch on the body. Beta activity reflects the mapping of touch in skin-based coordinates, whereas alpha activity reflects the mapping of touch in external space. We found that alpha activity was solely modulated by the location of tactile stimuli applied on a hand-held rod. Source reconstruction suggested that this alpha power modulation was localized in a network of fronto-parietal regions previously implicated in higher-order tactile and spatial processing. These findings are the first to implicate alpha oscillations in tool-extended sensing and suggest an important role for processing touch in external space when localizing touch on a tool.

## INTRODUCTION

Space is a crucial aspect of touch. Whether a mosquito lands on your arm, or a cat rubs against your leg, tactile events share the common feature of originating from somewhere, usually on the body. The sense of touch though, is not restricted to processing events on the body surface, as humans can also perceive tactile contact on external objects. For example, a blind person can use a white cane to locate obstacles in their immediate surroundings. Even in the sighted, when a person touches an object with the tip of a hand-held tool, the object is perceived to be at the tip instead of at the hand that holds the tool (Vaught et al., 1968; Yamamoto et al., 2005; Yamamoto and Kitazawa, 2001). Humans can also sense several of an object’s properties with a tool, such as its surface texture (Klatzky and Lederman, 1999; Yoshioka et al., 2007), softness (LaMotte, 2000), length (Peck et al., 1996) and position in space (Carello et al., 1992; Giudice et al., 2013).

Importantly, the ability to perceive touch on a tool extends along its entire surface, allowing humans to accurately localize where a hand-held tool is touched as they would on their own body (Miller et al., 2018). It was also found that *where* a tool is touched is likely rapidly encoded by location-specific patterns of vibration, termed *vibratory motifs*. Simulations with a biologically plausible skin-neuron model (Saal et al., 2017) suggested that these motifs are likely re-encoded by the spiking patterns of the hand’s mechanoreceptors, reflecting the initial transformation underlying the extraction of contact location on a rod. However, the nature of these mechanisms, particularly at the cortical level, is still largely unexplored. The aim of the present study is therefore to address the question of how the brain extracts ‘where’ a tool has been touched.

Since it is essential to identify the position of a tactile object in order to interact with it, more than a century of research has characterized the perceptual processes underlying tactile localization at a behavioral level ((Badde et al., 2015; Cholewiak and Collins, 2003; Fuchs et al., 2019; Harrar and Harris, 2009; Ho and Spence, 2007; Liu and Medina, 2021; Mancini et al., 2011; Miller et al., 2020; Parrish, 1897; Sadibolova et al., 2018; Stevens, 1992; Weber, 1846); for a review see (Heed et al., 2015)). These studies have identified different spatial codes that are utilized to localize tactile stimuli. The location of touch is thought to be initially encoded in skin-based coordinates—also called anatomical coordinates—and later integrated with extra-cutaneous information (e.g., proprioception) to remap its location in coordinates anchored to external space (Aglioti et al., 1999; Azañón and Soto-Faraco, 2008; Badde et al., 2015, 2014; Longo et al., 2015; Schicke and Roder, 2006; Shore et al., 2005, 2002). There has been extensive investigation on how these spatial codes are implemented at the neural level ((Celesia, 1979; Ghazanfar et al., 2000; Johansson and Flanagan, 2009; Johnson and Lamb, 1981; Tamè et al., 2015); for a review see (Dijkerman and de Haan, 2007)). Early activity in primary somatosensory cortex (SI) likely implements an anatomical code, as its layout is spatially organized in a limb-centric fashion (Penfield and Boldrey, 1937; Schnitzler et al., 1995). External coding, on the other hand, requires the involvement of a larger network that includes secondary somatosensory cortex (SII) and fronto-parietal regions (Azañón et al., 2010; Graziano and Gross, 1998; Reed et al., 2005; Soto-Faraco and Azañón, 2013; Takahashi et al., 2013; Wada et al., 2012).

Over the last decade, research using magnetoencephalography (MEG) and electroencephalography (EEG) has identified oscillatory mechanisms that underlie these processes. Touching the skin leads to a desynchronization in two main low-frequency bands, alpha (7-14 Hz) (Cheyne et al., 2003; Haegens et al., 2014; Neuper et al., 2006; Salenius et al., 1997; Salmelin and Hari, 1994) and beta (15-30 Hz) (Neuper et al., 2006; Pfurtscheller et al., 2001; Salmelin and Hari, 1994). Both frequency bands have been implicated in the spatial processing of touch (Heed et al., 2015). The alpha band has mainly been involved in the mapping of touch in the external coordinate system (Buchholz et al., 2013, 2011; Ruzzoli and Soto-Faraco, 2014; Schubert et al., 2019, 2015). Activity in the beta band, on the other hand, seems to exclusively reflect the mapping of touch in skin-centered coordinates (Buchholz et al., 2013, 2011b; Schubert et al., 2015). To date, whether these frequency bands also relate to spatial processing of touch on hand-held tools remains unknown. Our goal here was therefore to fill this gap.

We used EEG to investigate the oscillatory correlates of tool-extended tactile localization. Participants performed a delayed match-to-sample task that required them to judge the relative location of two successive contacts on a hand-held rod. Our previous event-related potentials (ERP) investigation (Miller et al., 2019) identified signatures of tactile localization starting as early as 48 ms after touch on a tool. Multivariate decoding algorithms and cortical source reconstruction suggested that the somatosensory cortex reuses low-level neural processes devoted to mapping touch on the body for mapping touch on the tool. Expanding on these previous results, we report here on the induced oscillations following touch localization on a tool. Given their role on mapping touch on the body, we focused our attention on the activity of the two aforementioned frequency bands: alpha and beta. Since tool use shares similar neural processes with body-related actions (Gallivan et al., 2013; Johnson-Frey, 2004) and based on our ERPs findings, we hypothesized that activity in both frequency bands would index spatial coding of touch on a hand-held tool.

## METHOD

Analyses were performed on a dataset for which we have previously analyzed ERPs following tactile stimulation (Miller et al., 2019). Here we focused on the induced oscillations and not oscillations phase-locked to the event.

### Participants

16 right-handed participants (mean age: 26.5 years, range 20 to 34 years, 5 males) free of any known sensory, perceptual, or motor disorders, volunteered to participate in the experiment. We chose this sample size in accordance to previous studies about somatosensory oscillations. The experiment was performed in accordance with the ethical standards laid down in the Declaration of Helsinki (2013) and all participants provided written informed consent according to national guidelines of the ethics committee (CPP SUD EST IV).

### Setup

Participants sat in a chair with their right arm placed on an adjustable armrest. During the task, they fixated on a central cross (2 cm wide) that was displayed on a 16’’ monitor ∼100 cm in front of them and aligned with their body midline. A one-meter long wooden rod was placed in their right hand, with the tip of the rod resting on a support so that it would stay stable and parallel to the participant’s body midline. Two solenoids (Mecalectro 8.19-.AB.83; 24 V, supplied with 36 W) were used to contact the rod at two different locations with a force of ∼15 N: ∼16 cm from the hand (close location) and ∼63 cm from the hand (far location). To ensure uniform and consistent contact between solenoids and the body of the rod, a plastic disc (4 cm diameter) was attached perpendicular to the tip of each solenoid’s metal rod. Participants used a double-pedal (Leptron Footswitch 548561) under their left foot to report their judgments.

In order to mask the sound generated by the solenoids, white noise was played continuously over noise-cancelling earphones (Bose QuietComfort 20) and a third solenoid was placed next to the rod between the other two solenoids; this ‘decoy’ solenoid was used to mask any residual auditory spatial cue to the location of impact. All solenoids were supported by adjustable tripods at the same height. Visual feedback was also prevented by spreading a black sheet between the screen and the participants’ neck to cover their hands and the stick.

### Experimental paradigm

Participants performed a delayed match-to-sample task designed to measure repetition effects on brain responses (Grill-Spector et al., 2006) during tactile localization on a tool (Figure 1B). In each trial, two hits were successively applied to the surface of the rod (Figure 1A). Participants had then to decide whether the second hit was in the same location (same close or far from the hand), or in a different location with respect to the first hit. Participants never wielded the rod before performing the task and never received feedback about their performance, ensuring that they could not learn arbitrary rules to distinguish close or far.

**Figure 1.**
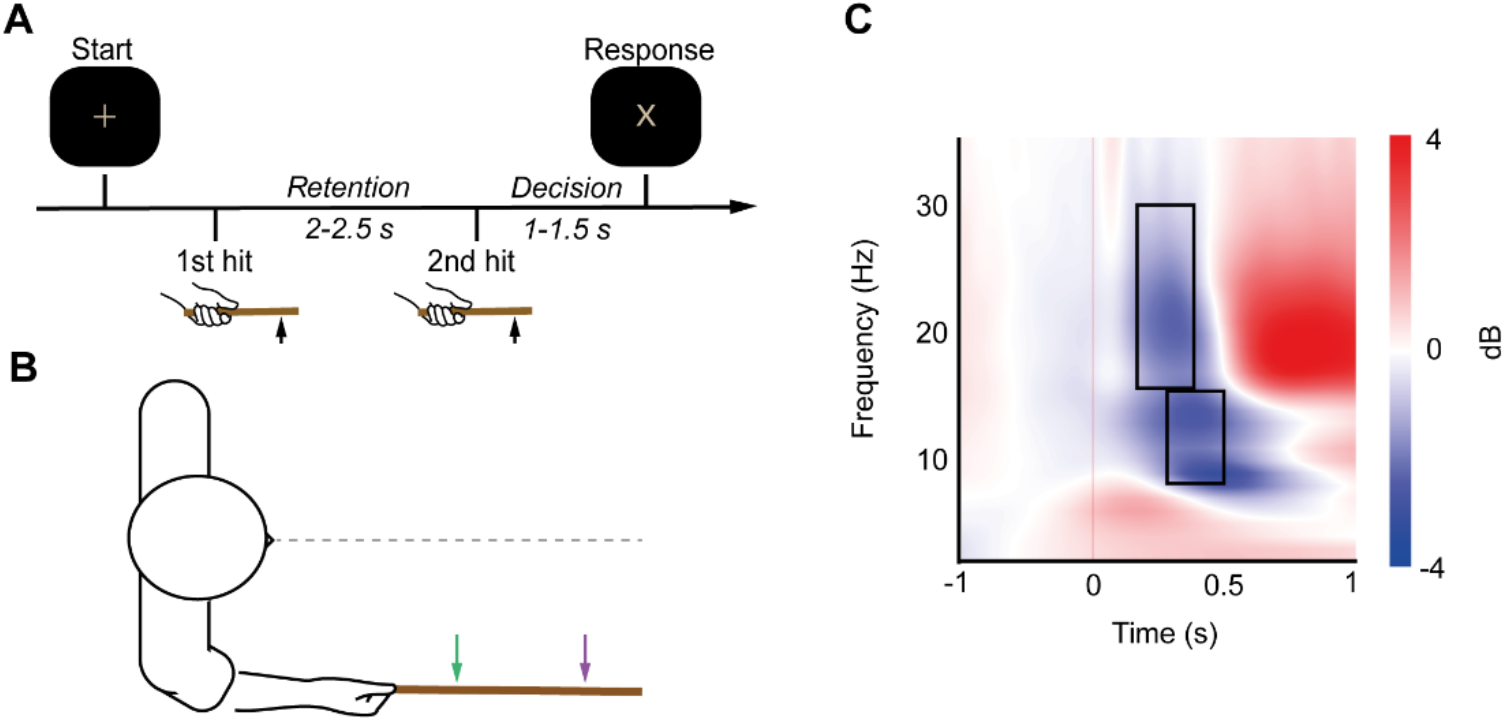
Experimental setup and paradigm. (A) Trial structure of the delayed match-to-sample task. For presentation purposes, only a portion of the rod is shown and black boxes represent the computer screen. (B) Participants (n=16) performed a 2-interval delayed match-to-sample task for touches applied at two locations (colored arrows) on the surface of a tool held in their right hand. This task forced participants to discriminate where the tool was touched. (C) Induced temporal dynamics of the after-hit period over contralateral somatosensory cortex. Oscillatory dynamics of C3 electrode for all epochs combined. Modulations are displayed as compared relative to baseline (−500 to -100 ms). Selected time windows for analysis are represented by grey rectangle for each frequency band: 300-500 ms for alpha and 200-400 ms for beta.

Each trial (Figure 1A) started when a fixation cross was displayed at the center of the screen. After 1000 ms, the fixation cross blinked to inform participants that the rod was about to be contacted. After a variable delay (between 1000–1500 ms; randomly chosen from a uniform distribution), a first stimulus was applied to the stick at one of the two locations (close or far; pseudorandomized), followed by a further delay (between 2000–2500 ms; randomly chosen from a uniform distribution) during which participants had to retain in memory the location of contact. The second stimulus (location chosen pseudorandomly) was then presented after which participants made a decision about whether the second hit was in the same or a different location as the first hit. They were instructed to wait until the fixation cross changed to an X (between 1000–1500 ms), prompting them to respond. The X turned back into a fixation cross just after the response was made, or after 2000 ms if no response. The next trial began after a fixed delay of 1000 ms.

Participants were instructed to discriminate whether the two contacts were in the same or a different location, using the foot pedals in the following way: if they judged that both hits were close to the hand they lifted their heel releasing the lower pedal. If they judged that both hits were far from the hand they lifted their toes releasing the upper pedal. If the two stimuli were different (close/far or far/close) they had to refrain from responding. To ensure that motor and perceptual processes wouldn’t overlap in the EEG data, participants used their left foot to make the response, thus engaging the sensorimotor system in the right hemisphere (i.e., ipsilateral to the stimulated hand). Participants performed four blocks of 100 trials each (400 trials in total), in which stimulus location was pseudorandomly interleaved for each hit. There were 200 contacts per location for each of the two hits by block (400 close and 400 far contacts in total). Furthermore, on half of the trials the two hits were in the same location and on the other half the two hits were in different locations. Each trial lasted between 5000 and 8500 ms. A brief rest period was provided between blocks during which participants could move their hands and eyes freely.

### EEG recording

EEG data were recorded continuously using a 65 channel ActiCap system (Brain Products). Horizontal and vertical electro-oculograms (EOGs) were recorded using electrodes placed below the left eye, and near the outer canthi of the right eye. Impedance of all electrodes was kept at <20 kΩ. FCz served as the online reference. EEG and EOG signals were low-pass filtered online at 0.1 Hz, sampled at 2500 Hz, and then saved to disk.

Stimulus presentation and behavioral response collection were performed using MatLab on the experimental control computer, which was synchronized and communicated with the EEG data recording system. The trial events were sent to the EEG data recording system via a parallel port.

### Pre-processing of the EEG data

EEG signals were preprocessed using the EEGLab Toolbox (Delorme and Makeig, 2004). The preprocessing steps for each participant were as follows: each participant’s four blocks of the experiment were appended into a single dataset. The signal was resampled at 250 Hz and high-pass filtered at 1 Hz. Faulty channels were interpolated using a spherical spline. We then epoched data into a time window of 3 seconds, 1 second before and 2 seconds after the second hit (time zero). Next, we removed signal artifacts with two steps: first, we removed eye blinks and horizontal eye movements from the signal using independent components analysis (ICA (Delorme and Makeig, 2004)) and a semi-automated algorithm called SASICA (Chaumon et al., 2015). Second, we excluded the first ten trials of the experiment, trials that were interrupted by the experimenter, all incorrectly answered trials, and manually rejected trials that were contaminated by muscle artifacts or other forms of signal noise. A response was determined as correct if: when the same two contacts happened, the participant responded for same, and gave the right location (far or close); when different contacts happened, the participant made no response. Only trials with correct response were kept for the analysis (mean accuracy = 96.4 ± 0.71%; range: 89.7%–99.5%). In total, this led to a mean exclusion of 35.5 trials per participant (range: 14–56). Next, we used the EEGLab function *pop_reref* to add FCz (the online reference) back into the dataset. Finally, we re-referenced the data to the average voltage across the scalp.

### Time-frequency decomposition

Time-frequency decomposition was performed using the open source toolbox Brainstorm (Tadel et al., 2011) in Matlab. The raw signal of each epoch was decomposed into frequencies between 1–35 Hz (linearly spaced) using Complex Morlet wavelets with a central frequency of 1 Hz and a full-width half maximum of 3 s. These parameters were chosen to ensure that our time-frequency decomposition had good spectral and temporal resolution within the chosen frequency range. The evoked activity was removed from the signal before time-frequency decomposition since it has already been analyzed in a previous study (Miller et al., 2019). Signal was then normalized in decibel (dB) to its ratio with the respective channel mean power during a baseline period ranging from -500 to -100 ms before hit. This period was free of any stimulus-induced activity that would extend into the baseline due to the temporal smoothing of the wavelet transformation, which is especially problematic for the lower frequencies (Cohen, 2014).

### EEG data analysis

Alpha- and beta-band activity were defined here as 7–14 Hz and 15–30 Hz respectively. Based on the observation of the oscillatory temporal dynamics during the time period after the hit (collapsed across all participants and conditions, see Figure 1C), we selected two time windows for analysis for each of our frequency band of interest: 200-400 ms for beta-band, and 300-500 ms for the alpha-band. This is a bias-free method for choosing time windows upon which running the analysis (Cohen, 2014).

We based our main analysis on repetition effects as a way of capturing correlates of localization devoid of potential low level confounds, such as intensity differences. Repeating a stimulus has been shown to modulate the power in frequency bands that index the processing of that feature (Grill-Spector et al., 2006). Thus, if the second hit occurred in the same location as the first hit, the power in frequency band(s) indexing spatial processing should significantly differ from when the second hit was in a different location. In order to reveal the modulation of activity related to contact location processing, we statistically compared activity when location of the second hit was repeated (in reference to the first hit of the same trial) versus when it was not repeated. To this aim, we averaged the power of each frequency band within the given time window, as well as the time points of each window, and compared the scalp topography of each condition (repeated, non-repeated) using a cluster-based permutation test (Maris and Oostenveld, 2007). Analysis was two-tailed with a cluster-level significance threshold set to 0.05 and 1000 permutations run.

### Source reconstruction

We performed source reconstruction for each epoch using the open source toolbox Brainstorm (Tadel et al., 2011). First, a head model was computed using OpenMEEG BEM model (Gramfort et al., 2010). A noise covariance matrix for every participant was computed over a baseline time window of -500 to -100 ms before stimulation. Sources were then estimated using the Standardized low resolution brain electromagnetic tomography (sLORETA (Pascual-Marqui, 2002)) approach with unconstrained dipole orientations across the surface.

We then performed time-frequency decomposition on the source files to localize significant power modulations in the alpha-band, since it returned significant location repetition effects. The signal at each vertex was decomposed into the mean of frequencies going from 7 to 14 Hz using Complex Morlet wavelets with a central frequency of 1 Hz and a full-width half maximum of 3 s. The signal was then normalized in decibel (dB) to its ratio with the respective channel mean power during a baseline period ranging from -500 to -100 ms before hit. A similar statistical comparison as the previous analysis was then applied in order to reveal the cerebral regions involved in processing contact location. We analyzed the difference in alpha power between each condition (repetition vs. non-repetition) during the same time windows (300 to 500 ms after hit) using a cluster-based comparison (Maris and Oostenveld, 2007).

## RESULTS

### Temporal dynamics of alpha and beta over contralateral somatosensory cortex

We first examined the temporal dynamics of oscillations at C3, an electrode directly above sensorimotor cortex (Oostenveld and Praamstra, 2001). Tactile stimulation on the rod led to a pattern of low-frequency power modulations (Figure 1C) over sensorimotor cortex that was consistent with prior work (Cheyne et al., 2003; Nierula et al., 2013). Specifically, we observed a desynchronization of alpha power between ∼250 to 600 ms after touch, and narrower desynchronization of beta power between 200 to 500 ms. Desynchronization within these time periods has been observed in previous studies (Neuper et al., 2006; Salmelin and Hari, 1994), including those related to spatial processing within each frequency band (Buchholz et al., 2013, 2011; Schubert et al., 2019, 2015).

Based on these changes in power spectra, analysis of alpha (Fig. 2) and beta (Fig. 3) was performed on distinct time windows that captured desynchronization of all of the frequencies in each band.

**Figure 2.**
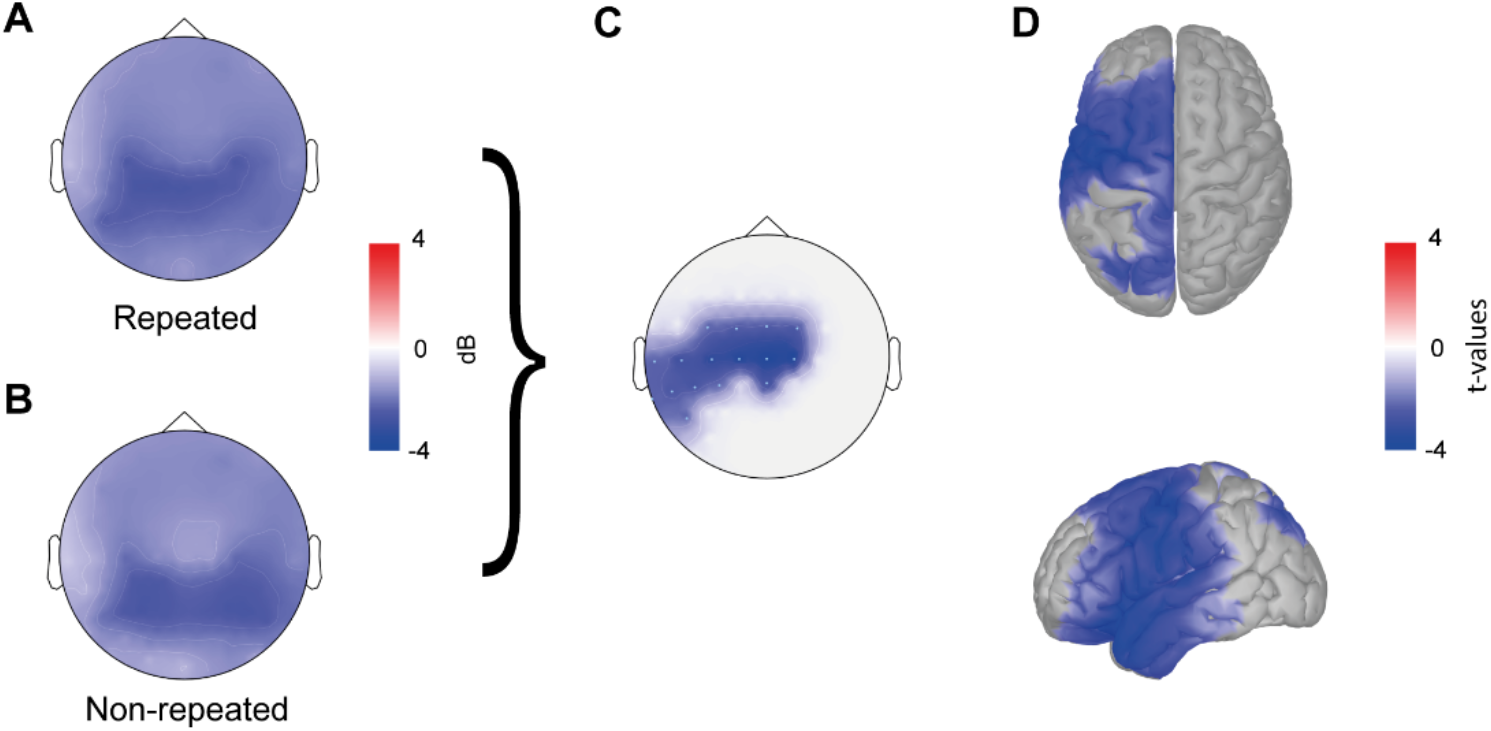
Alpha power modulation depends on contact location. (A-B) Average alpha-band (7–14 Hz) activity relative to baseline for the selected time windows (300-500 ms) following contact on the rod, when: A. location of the second hit was the same as the first hit and (B) location of the second hit was different than the 1^st^. (C) Scalp topography of the alpha power difference relative to contact location (Repeated vs.Non-repeated analysis, p = 0.024). (D) Source localization of the significant alpha power difference (p = 0.006). T-scale on the right is used for both (C) and (D).

**Figure 3.**
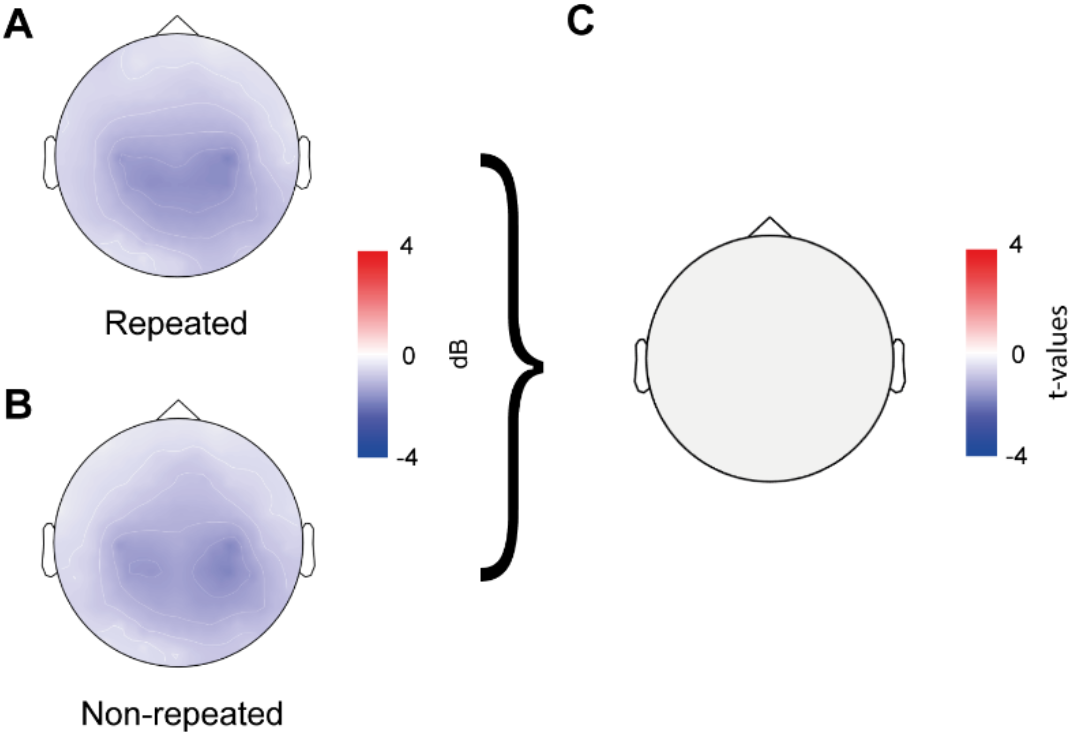
Beta power modulation doesn’t depend on contact location. (A-B) Average beta-band (15–30 Hz) activity relative to baseline for the selected time windows (200-400 ms) following contact on the rod, when: (A) location of the second hit was the same as the first hit and (B) location of the second hit was different than the first. (C) Scalp topography of the beta power difference relative to contact location (Repeated vs. Non-repeated analysis, no cluster found).

### Suppression in the alpha-band was dependent on location-based repetition

Touch on the rod led to widespread desynchronization in alpha power across the scalp, between 300 to 500 ms after touch. Consistent with previous studies (Buchholz et al., 2011; Schubert et al., 2019), this desynchronization was largely concentrated over centro-parietal channels bilaterally. As can be seen in Fig. 2A and B, the overall pattern of desynchronization on the scalp was largely independent of whether touch location was repeated or not (Figure 2A and B). Instead, location repetition modulated the gain of this desynchronization. We found significant differences in alpha power above central and parietal channels over the sensorimotor cortex contralateral to the stimulation (cluster-based p-value= 0.024). This difference was contralateral to the location of touch (Figure 2C), suggesting that the modulation observed is related to the tactile location processing and not to the motor preparation of the response.

We followed up by identifying the cortical sources underlying the observed alpha modulation. We found widespread repetition effects in alpha throughout the frontal, parietal, and temporal cortices (Figure 2D, p = 0.006). These sources notably include SI, SII, dorsal and ventral premotor cortex, occipito-parietal cortex, and the superior temporal gyrus (STG).

### Beta-band activity was independent from location-based repetition

We observed a desynchronization of beta power over bilateral centro-parietal channels between 200 to 400 ms after touch on the rod. As can be seen in Figure 3A and B, whether location was repeated or not did not affect the scalp distribution of desynchronization. Indeed, a cluster-based permutation test for the selected time window revealed no significant difference in beta power (Figure 3C). Therefore, in contrast to the alpha band and with our initial hypothesis, we did not observe repetition effects in the beta band.

## DISCUSSION

The present study aimed to investigate whether oscillatory activity linked to spatial coding of touch on the body (Buchholz et al., 2013, 2011; Ruzzoli and Soto-Faraco, 2014; Schubert et al., 2018, 2015) is also involved in sensing location through a hand-held tool. To this end, we compared oscillatory EEG responses in the alpha (7–14 Hz) and beta (15–30 Hz) bands during a delayed match-to-sample task where two successive touches were applied to either the same or different location on a hand-held wooden rod. Contact location information was isolated based on repetition effects, by comparing power modulation when location was repeated to when it was not.

We found that the post-stimulus desynchronization of alpha activity between 300 to 500 ms after touch was differentially affected by whether contact location was repeated. In contrast, we did not observe any repetition effects in the post-stimulus desynchronization of beta activity. Moreover, source estimation of the observed alpha modulation showed it originated in a contralateral frontoparietal network that is known to be involved in tactile localization. These findings provide novel evidence for the involvement of alpha-band activity in spatial coding of touch on a tool.

Owing to the remarkable accuracy with which humans can similarly localize touch on their arm and on a hand-held tool, previous work hypothesized that similar mechanisms may be involved in both (Miller et al., 2018; Yamamoto et al., 2005). In a recent event-related potentials study, we provided evidence that location information encoded in the tool’s vibratory pattern is rapidly extracted and processed (<100 ms post-touch) by somatosensory and parietal cortices. Furthermore, these location-related neural responses were similar to those observed when touch was on the arm; this suggests that the brain may reuse neural mechanisms dedicated to processing touch on the body to process touch on a tool (Miller et al., 2019).

The present study focused on assessing whether alpha and beta oscillations—known to be associated to localizing touch on the body in external and anatomical reference frames, respectively (Buchholz et al., 2013, 2011; Schubert et al., 2019, 2015)—are also involved in localizing touch on a tool. In keeping with previous reports, we observed a post-touch desynchronization in both alpha and beta power when touch was applied to a hand-held tool. In addition, their temporal distributions were consistent with what has been previously shown for touch on the skin (Cheyne et al., 2003; Gaetz and Cheyne, 2006). Importantly, repeating contact location on the tool allowed us to isolate oscillatory activity specific to spatial processing. That is, a repetition effect in oscillatory power served as a signature of localization-related processing. We identified such a signature in the alpha-band activity, with a significant effect of repetition on desynchronization in the hemisphere contralateral to the stimulation (Figure 2A-C). In contrast, the magnitude of the beta desynchronization was not modulated by whether touch location on the tool was repeated or not (Figure 3). This suggests that only the alpha-band activity is involved in localization of touch on a tool.

These findings raise the question as to which aspects of sensorimotor processing are reflected in the observed alpha desynchronization. One possibility, which we can readily rule out, is that it reflects motor preparatory activity. Such activity would likely be observed contralateral to the foot used for the response (i.e., the left) and therefore right lateralized (i.e., ipsilateral to the stimulation). However, the repetition effects in both the sensor-level activity (Figure 2A-C) and the sources (Figure 2D) were all lateralized to the left hemisphere (i.e., contralateral to stimulation), underscoring its relation to the processing of tactile information and not motor preparation.

The repetition effects in alpha-band activity instead likely reflect the processing of spatial tactile information on the surface of a hand-held tool. As previously discussed, alpha activity has been implicated in tactile spatial processing, being thought to play a prominent role in the coding of external spatial information for touch (Buchholz et al., 2013, 2011; Ruzzoli and Soto-Faraco, 2014; Schubert et al., 2019). Our findings provide initial evidence that alpha activity is also involved in tactile localization on a tool, thus possibly implicating the use of an external reference frame for the coding of tactile information on tools.

Our source localization of the repetition effect in the alpha-band activity is consistent with this proposition. Repeating the location of touch modulated activity in a left-lateralized cortical network that included motor and somatosensory cortices, premotor (PMC) and posterior parietal cortices (PPC), and temporal regions (Figure 2D). Studies in both human and non-human primates have implicated these regions—particularly the PMC and PPC—in processing touch in external space (Avillac et al., 2005; Azañón et al., 2010; Bremmer et al., 2001; Lloyd et al., 2003). Furthermore, alpha oscillatory activity in these regions appears to play a crucial role in this process (Buchholz et al., 2011; Ruzzoli and Soto-Faraco, 2014; Schubert et al., 2019).

Post-touch alpha oscillations may reflect the orienting of attention in external space (Ossandón et al., 2020), which is involved in the alignment of sensory modalities under a common coordinate system (Ruzzoli and Soto-Faraco, 2014). Indeed, the PMC and PPC are crucial for the reference frame transformations underlying multimodal integration (Avillac et al., 2005; Bolognini and Maravita, 2007; Bremmer et al., 2001; Graziano and Gross, 1998). Vallar and Maravita have argued that both regions are also important for making the experience of space integrated and unitary across modalities (Vallar and Maravita, 2009). Consistent with this proposal, alpha-band activity has also been implicated in spatial processing of visual (Foster et al., 2017, 2016; Rihs et al., 2007; Samaha et al., 2016) and auditory (Deng et al., 2020; Frey et al., 2014) stimuli. In the context of these findings, the observed alpha desynchronization during tool extended sensing might reflect a supramodal mechanism for spatial localization.

Overall, while our findings are clear in suggesting a role of alpha oscillations in localizing touch on a hand-held tool, the nature of the spatial representation remains unclear. The present study was not aimed at disentangling the actual coordinate system reflected in tool-extended sensing. Indeed, isolating tactile reference frames typically requires manipulating the posture of an effector, such as crossing the hands. When the hands are crossed, anatomical (e.g. right hand) and external (e.g. left side) reference frames can be spatially dissociated. Thus, to precisely pinpoint the spatial code(s) used in tool-extended tactile localization, future studies are needed that manipulate tool posture in a similar way, possibly allowing to dissociate the signatures of anatomical and external reference frames.

## CONCLUSION

To conclude, we newly report that alpha activity is specifically modulated by the location of tactile stimuli applied on a hand-held rod. The neural sources of this alpha power modulation were localized to a network of parieto-frontal areas previously implicated in tactile and spatial processing. While further studies are needed to deepen the specific role of brain oscillations in processing touch on tools, we provide novel evidence involving alpha band activity in tool-extended sensing.

## DATA AVAILABILITY

All data will be available in an online repository upon acceptance.

## AKNOWLEDGEMENTS

We thank Valeria Ravenda for helping collect the data and Frederic Volland for his help constructing the experimental setup.

## AUTHOR CONTRIBUTIONS

CF, LEM and AF conceived of the experimental idea, designed the EEG experiment, and wrote the paper. CF and LEM collected and analyzed the data. RS and EK built the solenoid setup. All authors approved of the final submission.

## REFERENCES

Aglioti, S., Smania, N., Peru, A., 1999. Frames of Reference for Mapping Tactile Stimuli in Brain-Damaged Patients. Journal of Cognitive Neuroscience 11, 67–79. https://doi.org/10.1162/089892999563256

Avillac, M., Denève, S., Olivier, E., Pouget, A., Duhamel, J.-R., 2005. Reference frames for representing visual and tactile locations in parietal cortex. Nature Neuroscience 8, 941–949. https://doi.org/10.1038/nn1480

Azañón, E., Longo, M.R., Soto-Faraco, S., Haggard, P., 2010. The Posterior Parietal Cortex Remaps Touch into External Space. Current Biology 20, 1304–1309. https://doi.org/10.1016/j.cub.2010.05.063

Azañón, E., Soto-Faraco, S., 2008. Changing Reference Frames during the Encoding of Tactile Events. Current Biology 18, 1044–1049. https://doi.org/10.1016/j.cub.2008.06.045

Badde, S., Röder, B., Heed, T., 2015. Flexibly weighted integration of tactile reference frames. Neuropsychologia 70, 367–374. https://doi.org/10.1016/j.neuropsychologia.2014.10.001

Badde, S., Röder, B., Heed, T., 2014. Multiple spatial representations determine touch localization on the fingers. Journal of Experimental Psychology: Human Perception and Performance 40, 784–801. https://doi.org/10.1037/a0034690

Bolognini, N., Maravita, A., 2007. Proprioceptive Alignment of Visual and Somatosensory Maps in the Posterior Parietal Cortex. Current Biology 17, 1890–1895. https://doi.org/10.1016/j.cub.2007.09.057

Bremmer, F., Schlack, A., Duhamel, J.-R., Graf, W., Fink, G.R., 2001. Space Coding in Primate Posterior Parietal Cortex. NeuroImage 14, S46–S51. https://doi.org/10.1006/nimg.2001.0817

Buchholz, V.N., Jensen, O., Medendorp, W.P., 2013. Parietal Oscillations Code Nonvisual Reach Targets Relative to Gaze and Body. Journal of Neuroscience 33, 3492–3499. https://doi.org/10.1523/JNEUROSCI.3208-12.2013

Buchholz, V.N., Jensen, O., Medendorp, W.P., 2011a. Multiple Reference Frames in Cortical Oscillatory Activity during Tactile Remapping for Saccades. Journal of Neuroscience 31, 16864–16871. https://doi.org/10.1523/JNEUROSCI.3404-11.2011

Buchholz, V.N., Jensen, O., Medendorp, W.P., 2011b. Multiple Reference Frames in Cortical Oscillatory Activity during Tactile Remapping for Saccades. Journal of Neuroscience 31, 16864–16871. https://doi.org/10.1523/JNEUROSCI.3404-11.2011

Carello, C., Fitzpatrick, P., Turvey, M.T., 1992. Haptic probing: Perceiving the length of a probe and the distance of a surface probed. Perception & Psychophysics 51, 580–598. https://doi.org/10.3758/BF03211655

Celesia, G.G., 1979. Somatosensory Evoked Potentials Recorded Directly From Human Thalamus and Sm I Cortical Area. Archives of Neurology 36, 399–405. https://doi.org/10.1001/ar-chneur.1979.00500430029003

Chaumon, M., Bishop, D.V.M., Busch, N.A., 2015. A practical guide to the selection of independent components of the electroencephalogram for artifact correction. Journal of Neuroscience Methods 250, 47–63. https://doi.org/10.1016/j.jneumeth.2015.02.025

Cheyne, D., Gaetz, W., Garnero, L., Lachaux, J.-P., Ducorps, A., Schwartz, D., Varela, F.J., 2003. Neuromagnetic imaging of cortical oscillations accompanying tactile stimulation. Cognitive Brain Research 17, 599–611. https://doi.org/10.1016/S0926-6410(03)00173-3

Cholewiak, R.W., Collins, A.A., 2003. Vibrotactile localization on the arm: Effects of place, space, and age. Perception & Psychophysics 65, 1058–1077. https://doi.org/10.3758/BF03194834

Cohen, M.X., 2014. Analyzing Neural Time Series Data: Theory and Practice. MIT Press.

Delorme, A., Makeig, S., 2004. EEGLAB: an open source toolbox for analysis of single-trial EEG dynamics including independent component analysis. Journal of Neuroscience Methods 134, 9– 21. https://doi.org/10.1016/j.jneumeth.2003.10.009

Deng, Y., Choi, I., Shinn-Cunningham, B., 2020. Topographic specificity of alpha power during auditory spatial attention. NeuroImage 207, 116360. https://doi.org/10.1016/j.neuroimage.2019.116360

Dijkerman, H.C., de Haan, E.H.F., 2007. Somatosensory processes subserving perception and action. Behavioral and Brain Sciences 30, 189. https://doi.org/10.1017/S0140525X07001392

Foster, J.J., Bsales, E.M., Jaffe, R.J., Awh, E., 2017. Alpha-Band Activity Reveals Spontaneous Representations of Spatial Position in Visual Working Memory. Current Biology 27, 3216-3223.e6. https://doi.org/10.1016/j.cub.2017.09.031

Foster, J.J., Sutterer, D.W., Serences, J.T., Vogel, E.K., Awh, E., 2016. The topography of alpha-band activity tracks the content of spatial working memory. Journal of Neurophysiology 115, 168– 177. https://doi.org/10.1152/jn.00860.2015

Frey, J.N., Mainy, N., Lachaux, J.-P., Muller, N., Bertrand, O., Weisz, N., 2014. Selective Modulation of Auditory Cortical Alpha Activity in an Audiovisual Spatial Attention Task. Journal of Neuroscience 34, 6634–6639. https://doi.org/10.1523/JNEUROSCI.4813-13.2014

Fuchs, X., Wulff, D.U., Heed, T., 2019. Online sensory feedback during active search improves tactile localization (preprint). Neuroscience. https://doi.org/10.1101/590539

Gaetz, W., Cheyne, D., 2006. Localization of sensorimotor cortical rhythms induced by tactile stimulation using spatially filtered MEG. NeuroImage 30, 899–908. https://doi.org/10.1016/j.neuroimage.2005.10.009

Gallivan, J.P., McLean, D.A., Valyear, K.F., Culham, J.C., 2013. Decoding the neural mechanisms of human tool use. eLife 2, e00425. https://doi.org/10.7554/eLife.00425

Ghazanfar, A.A., Stambaugh, C.R., Nicolelis, M.A.L., 2000. Encoding of Tactile Stimulus Location by Somatosensory Thalamocortical Ensembles. J. Neurosci. 20, 3761–3775. https://doi.org/10.1523/JNEUROSCI.20-10-03761.2000

Giudice, N.A., Klatzky, R.L., Bennett, C.R., Loomis, J.M., 2013. Perception of 3-D location based on vision, touch, and extended touch. Exp Brain Res 224, 141–153. https://doi.org/10.1007/s00221-012-3295-1

Gramfort, A., Papadopoulo, T., Olivi, E., Clerc, M., 2010. OpenMEEG: opensource software for quasistatic bioelectromagnetics 20.

Graziano, M.S., Gross, C.G., 1998. Spatial maps for the control of movement. Current Opinion in Neurobiology 8, 195–201. https://doi.org/10.1016/S0959-4388(98)80140-2

Grill-Spector, K., Henson, R., Martin, A., 2006. Repetition and the brain: neural models of stimulus-specific effects. Trends in Cognitive Sciences 10, 14–23. https://doi.org/10.1016/j.tics.2005.11.006

Haegens, S., Vazquez, Y., Zainos, A., Alvarez, M., Jensen, O., Romo, R., 2014. Thalamocortical rhythms during a vibrotactile detection task. Proceedings of the National Academy of Sciences 111, E1797–E1805. https://doi.org/10.1073/pnas.1405516111

Harrar, V., Harris, L.R., 2009. Eye position affects the perceived location of touch. Exp Brain Res 198, 403–410. https://doi.org/10.1007/s00221-009-1884-4

Heed, T., Buchholz, V.N., Engel, A.K., Röder, B., 2015. Tactile remapping: from coordinate transformation to integration in sensorimotor processing. Trends in Cognitive Sciences 19, 251–258. https://doi.org/10.1016/j.tics.2015.03.001

Ho, C., Spence, C., 2007. Head orientation biases tactile localization. Brain Research 1144, 136–141. https://doi.org/10.1016/j.brainres.2007.01.091

Johansson, R.S., Flanagan, J.R., 2009. Coding and use of tactile signals from the fingertips in object manipulation tasks. Nat Rev Neurosci 10, 345–359. https://doi.org/10.1038/nrn2621

Johnson, K.O., Lamb, G.D., 1981. Neural mechanisms of spatial tactile discrimination: neural patterns evoked by braille-like dot patterns in the monkey. The Journal of Physiology 310, 117–144. https://doi.org/10.1113/jphysiol.1981.sp013540

Johnson-Frey, S.H., 2004. The neural bases of complex tool use in humans. Trends in Cognitive Sciences 8, 71–78. https://doi.org/10.1016/j.tics.2003.12.002

Klatzky, R.L., Lederman, S.J., 1999. Tactile roughness perception with a rigid link interposed between skin and surface. Perception & Psychophysics 61, 591–607. https://doi.org/10.3758/BF03205532

LaMotte, R.H., 2000. Softness Discrimination With a Tool. Journal of Neurophysiology 83, 1777–1786. https://doi.org/10.1152/jn.2000.83.4.1777

Liu, Y., Medina, J., 2021. Visuoproprioceptive conflict in hand position biases tactile localization on the hand surface. Journal of Experimental Psychology: Human Perception and Performance 47, 344–356. https://doi.org/10.1037/xhp0000893

Lloyd, D.M., Shore, D.I., Spence, C., Calvert, G.A., 2003. Multisensory representation of limb position in human premotor cortex. Nature Neuroscience 6, 17–18. https://doi.org/10.1038/nn991

Longo, M.R., Mancini, F., Haggard, P., 2015. Implicit body representations and tactile spatial remapping. Acta Psychologica 160, 77–87. https://doi.org/10.1016/j.actpsy.2015.07.002

Mancini, F., Longo, M.R., Iannetti, G.D., Haggard, P., 2011. A supramodal representation of the body surface. Neuropsychologia 49, 1194–1201. https://doi.org/10.1016/j.neuropsychologia.2010.12.040

Maris, E., Oostenveld, R., 2007. Nonparametric statistical testing of EEG- and MEG-data. Journal of Neuroscience Methods 164, 177–190. https://doi.org/10.1016/j.jneumeth.2007.03.024

Miller, L.E., Fabio, C., Ravenda, V., Bahmad, S., Koun, E., Salemme, R., Luauté, J., Bolognini, N., Hayward, V., Farnè, A., 2019. Somatosensory Cortex Efficiently Processes Touch Located Beyond the Body. Current Biology S0960982219313831. https://doi.org/10.1016/j.cub.2019.10.043

Miller, L.E., Fabio, C., van Beers, R., Farnè, A., Medendorp, W.P., 2020. A neural surveyor in somatosensory cortex (preprint). Neuroscience. https://doi.org/10.1101/2020.06.26.173419

Miller, L.E., Montroni, L., Koun, E., Salemme, R., Hayward, V., Farnè, A., 2018. Sensing with tools extends somatosensory processing beyond the body. Nature 561, 239–242. https://doi.org/10.1038/s41586-018-0460-0

Neuper, C., Wörtz, M., Pfurtscheller, G., 2006. ERD/ERS patterns reflecting sensorimotor activation and deactivation, in: Progress in Brain Research. Elsevier, pp. 211–222. https://doi.org/10.1016/S0079-6123(06)59014-4

Nierula, B., Hohlefeld, F.U., Curio, G., Nikulin, V.V., 2013. No somatotopy of sensorimotor alpha-oscillation responses to differential finger stimulation. NeuroImage 76, 294–303. https://doi.org/10.1016/j.neuroimage.2013.03.025

Oostenveld, R., Praamstra, P., 2001. The five percent electrode system for high-resolution EEG and ERP measurements. Clinical Neurophysiology 7.

Ossandón, J.P., König, P., Heed, T., 2020. No Evidence for a Role of Spatially Modulated α-Band Activity in Tactile Remapping and Short-Latency, Overt Orienting Behavior. J. Neurosci. 40, 9088–9102. https://doi.org/10.1523/JNEUROSCI.0581-19.2020

Parrish, C.S., 1897. Localisation of Cutaneous Impressions by Arm Movement without Pressure upon the Skin. The American Journal of Psychology 8, 250. https://doi.org/10.2307/1410941

Pascual-Marqui, R.D., 2002. Standardized low resolution brain electromagnetic. Clinical Pharmacology 16.

Peck, A.J., Jeffers, R.G., Carello, C., Turvey, M.T., 1996. Haptically Perceiving the Length of One Rod by Means of Another. Ecological Psychology 8, 237–258. https://doi.org/10.1207/s15326969eco0803_3

Penfield, W., Boldrey, E., 1937. SOMATIC MOTOR AND SENSORY REPRESENTATION IN THE CEREBRAL CORTEX OF MAN AS STUDIED BY ELECTRICAL STIMULATION. Brain 60, 389–443. https://doi.org/10.1093/brain/60.4.389

Pfurtscheller, G., Woertz, M., Krausz, G., Neuper, C., 2001. Distinction of different ®ngers by the frequency of stimulus induced beta oscillations in the human EEG. Neuroscience Letters 4.

Reed, C.L., Klatzky, R.L., Halgren, E., 2005. What vs. where in touch: an fMRI study. NeuroImage 25, 718–726. https://doi.org/10.1016/j.neuroimage.2004.11.044

Rihs, T.A., Michel, C.M., Thut, G., 2007. Mechanisms of selective inhibition in visual spatial attention are indexed by ?-band EEG synchronization. Eur J Neurosci 25, 603–610. https://doi.org/10.1111/j.1460-9568.2007.05278.x

Ruzzoli, M., Soto-Faraco, S., 2014. Alpha Stimulation of the Human Parietal Cortex Attunes Tactile Perception to External Space. Current Biology 24, 329–332. https://doi.org/10.1016/j.cub.2013.12.029

Saal, H.P., Delhaye, B.P., Rayhaun, B.C., Bensmaia, S.J., 2017. Simulating tactile signals from the whole hand with millisecond precision. Proceedings of the National Academy of Sciences 114, E5693–E5702. https://doi.org/10.1073/pnas.1704856114

Sadibolova, R., Tamè, L., Longo, M.R., 2018. More than skin-deep: Integration of skin-based and musculoskeletal reference frames in localization of touch. Journal of Experimental Psychology: Human Perception and Performance 44, 1672–1682. https://doi.org/10.1037/xhp0000562

Salenius, S., Schnitzler, A., Salmelin, R., Jousmäki, V., Hari, R., 1997. Modulation of Human Cortical Rolandic Rhythms during Natural Sensorimotor Tasks. NeuroImage 5, 221–228. https://doi.org/10.1006/nimg.1997.0261

Salmelin, R., Hari, R., 1994. Spatiotemporal characteristics of sensorimotor neuromagnetic rhythms related to thumb movement. Neuroscience 60, 537–550. https://doi.org/10.1016/0306-4522(94)90263-1

Samaha, J., Sprague, T.C., Postle, B.R., 2016. Decoding and Reconstructing the Focus of Spatial Attention from the Topography of Alpha-band Oscillations. Journal of Cognitive Neuroscience 28, 1090–1097. https://doi.org/10.1162/jocn_a_00955

Schicke, T., Roder, B., 2006. Spatial remapping of touch: Confusion of perceived stimulus order across hand and foot. Proceedings of the National Academy of Sciences 103, 11808–11813. https://doi.org/10.1073/pnas.0601486103

Schnitzler, A., Salmelin, R., Salenius, S., Jousmäki, V., Hari, R., 1995. Tactile information from the human hand reaches the ipsilateral primary somatosensory cortex. Neuroscience Letters 200, 25–28. https://doi.org/10.1016/0304-3940(95)12065-C

Schubert, J.T.W., Buchholz, V.N., Föcker, J., Engel, A.K., Röder, B., Heed, T., 2019. Alpha-band oscillations reflect external spatial coding for tactile stimuli in sighted, but not in congenitally blind humans. Sci Rep 9, 9215. https://doi.org/10.1038/s41598-019-45634-w

Schubert, J.T.W., Buchholz, V.N., Föcker, J., Engel, A.K., Röder, B., Heed, T., 2015. Oscillatory activity reflects differential use of spatial reference frames by sighted and blind individuals in tactile attention. NeuroImage 117, 417–428. https://doi.org/10.1016/j.neuroimage.2015.05.068

Schubert, J.T.W., Buchholz, V.N., Foecker, J., Engel, A.K., Roder, B., Heed, T., 2018. Alpha-band oscillations reflect external spatial coding for tactile stimuli in sighted, but not in congenitally blind humans. bioRxiv. https://doi.org/10.1101/442384

Shore, D.I., Gray, K., Spry, E., Spence, C., 2005. Spatial Modulation of Tactile Temporal-Order Judgments. Perception 34, 1251–1262. https://doi.org/10.1068/p3313

Shore, D.I., Spry, E., Spence, C., 2002. Confusing the mind by crossing the hands. Cognitive Brain Research 14, 153–163. https://doi.org/10.1016/S0926-6410(02)00070-8

Soto-Faraco, S., Azañón, E., 2013. Electrophysiological correlates of tactile remapping. Neuropsychologia 51, 1584–1594. https://doi.org/10.1016/j.neuropsychologia.2013.04.012

Stevens, J.C., 1992. Aging and Spatial Acuity of Touch. Journal of Gerontology 47, P35–P40. https://doi.org/10.1093/geronj/47.1.P35

Tadel, F., Baillet, S., Mosher, J.C., Pantazis, D., Leahy, R.M., 2011. Brainstorm: A User-Friendly Application for MEG/EEG Analysis. Computational Intelligence and Neuroscience 2011, 1–13. https://doi.org/10.1155/2011/879716

Takahashi, T., Kansaku, K., Wada, M., Shibuya, S., Kitazawa, S., 2013. Neural Correlates of Tactile Temporal-Order Judgment in Humans: an fMRI Study. Cerebral Cortex 23, 1952–1964. https://doi.org/10.1093/cercor/bhs179

Tamè, L., Pavani, F., Braun, C., Salemme, R., Farnè, A., Reilly, K.T., 2015. Somatotopy and temporal dynamics of sensorimotor interactions: evidence from double afferent inhibition. Eur J Neurosci 41, 1459–1465. https://doi.org/10.1111/ejn.12890

Vallar, G., Maravita, A., 2009. Personal and Extrapersonal Spatial Perception, in: Berntson, G.G., Cacioppo, J.T. (Eds.), Handbook of Neuroscience for the Behavioral Sciences. John Wiley & Sons, Inc., Hoboken, NJ, USA, p. neubb001016. https://doi.org/10.1002/9780470478509.neubb001016

Vaught, G.M., Simpson, W.E., Ryder, R., 1968. Feeling with a stick. Perceptual and Motor Skills 26, 848–848. https://doi.org/10.2466/pms.1968.26.3.848

Wada, M., Takano, K., Ikegami, S., Ora, H., Spence, C., Kansaku, K., 2012. Spatio-Temporal Updating in the Left Posterior Parietal Cortex. PLoS ONE 7, e39800. https://doi.org/10.1371/journal.pone.0039800

Weber, E., 1846. Tastsinn und Gemeingefuhl [On the sense of touch and ‘common sensibility’]. E. H. Weber on the Tactile Senses, Earlbaum,. Taylor & Francis, Hove, UK.

Yamamoto, S., Kitazawa, S., 2001. Sensation at the tips of invisible tools. Nature neuroscience 4, 979–980.

Yamamoto, S., Moizumi, S., Kitazawa, S., 2005. Referral of Tactile Sensation to the Tips of L-Shaped Sticks. Journal of Neurophysiology 93, 2856–2863. https://doi.org/10.1152/jn.01015.2004

Yoshioka, T., Bensmaïa, S.J., Craig, J.C., Hsiao, S.S., 2007. Texture perception through direct and indirect touch: An analysis of perceptual space for tactile textures in two modes of exploration. Somatosensory & Motor Research 24, 53–70. https://doi.org/10.1080/08990220701318163

